# An open platform for high-resolution light-based control of microscopic collectives

**DOI:** 10.1101/2020.12.28.424547

**Authors:** Ana Rubio Denniss, Thomas E. Gorochowski, Sabine Hauert

## Abstract

Engineering microscopic collectives of cells or microrobots is challenging due to the often-limited capabilities of the individual agents, our inability to program their motion and local interactions, and difficulties visualising their behaviours. Here, we present a low-cost, modular and open-source Dynamic Optical MicroEnvironment (DOME) and demonstrate its ability to augment microagent capabilities and control collective behaviours using light. The DOME offers an accessible means to study complex multicellular phenomena and implement *de-novo* microswarms with desired functionalities.

## Main text

Self-organisation is used by many biological systems to drive the emergence of robust collective phenomena across large populations of cells and is known to underpin processes such as tissue morphogenesis, collective motion and disease progression ^1^. Engineering microagents – be they living cells or human-made microrobots – with similarly complex collective behaviours would have applications ranging from the design of new functional materials ^2^ to novel biomedical therapies ^3,4^. The key component driving self-organisation is the ability of agents to react to their local environment following simple behavioural rules ^5^. However, at present we struggle to rapidly tweak the rules microagents follow. As a stepping stone, we propose to externally control each microagent and their reaction to the local environment, allowing for the rapid prototyping of behavioural rules that give rise to self-organisation. Light is perfectly suited for this task and at small scales can be used to make and break bonds ^6^, power micromotors ^7^, alter shapes ^8^, drive the release of a cargo^9^, modify microenvironments ^10^, and interact with light sensing organisms ^11,12^, making it a powerful tool for microagent control. Unlike other methods based on the use of chemicals or magnetic fields, light is better suited to the simultaneous control of many agents due to its high spatio-temporal resolution.

Light-controlled microswarms have been shown to form sophisticated shapes ^13^, flock ^14^, perform collective phototaxis ^15^, self-assemble into active materials ^16^, and treat tumours ^17^, yet many of these systems rely on manual control of a single or few light stimuli, offering limited local control at the scale of large collectives. Where automated closed-loop high-resolution spatio-temporal control of microswarms is used ^13^, bespoke equipment is needed making the work inaccessible for most labs.

To address these limitations, here we present the Dynamic Optical MicroEnvironment (DONE), which is able to project dynamic light patterns in response to the behaviour of light-reactive microagents and guide their collective behaviour (**Figure 1a**). This is made possible by the DOME’s ability to continuously image a microsystem and use this information as input into feedback control schemes that can then modify the light pattern projected in real-time to interact and guide the behaviours of the individual agents. Furthermore, the DOME has been specifically designed to be low-cost, modular, and open source to allow for the easy adaptation to new applications. This builds on other exciting open platforms focussed on microscopy, by adding fine light-based control ^18^.

**Figure 1:**
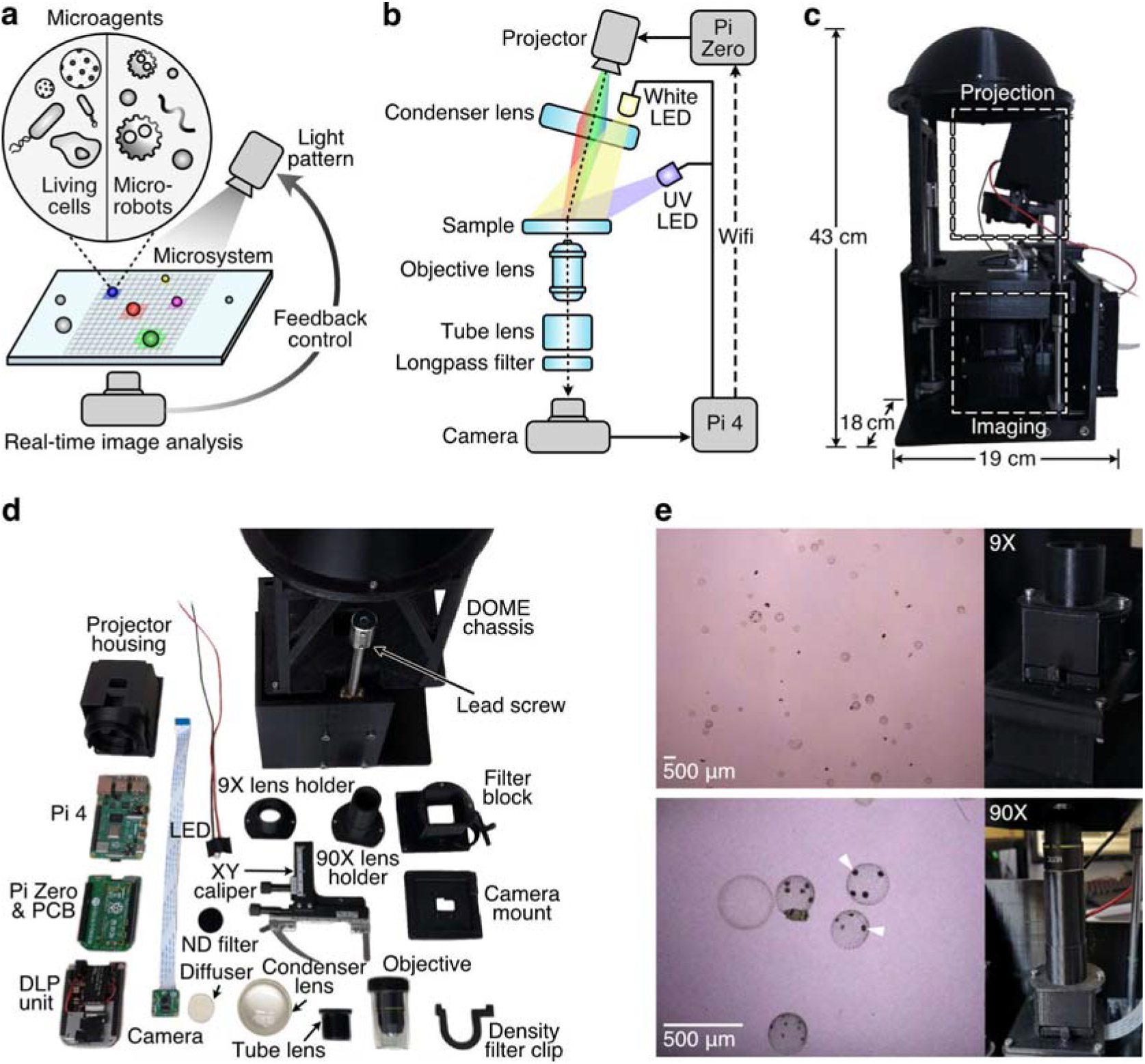
The Dynamic Optical MicroEnvironment (DOME). (**a**) Conceptual diagram of using light-based feedback control for diverse microagent collectives. (**b**) Schematic of key components and their connection within the DOME. Optical components are coloured light-blue, and arrows depict the flow of information. Dotted lines denote the optical path and dashed lines are wireless communication links. (**c**) Fully assembled DOME with dimensions. (**d**) Dismantled DOME with all key components labelled. (**e**) *Volvox* colonies imaged through a 9X magnification lens (top) and 90X magnification lens (bottom). White arrows show daughter *Volvox* colonies. Images on the left show the optical setup for each configuration.

The DOME consists of three major subsystems: dynamic light projection, real-time imaging, and generation of feedback control signals (**Figure 1b–d**). Light projection is performed by a digital light projector that creates a controllable grid of 854 × 480 pixels (0.4 megapixels) where each pixel can have its red, blue, and green (RGB) intensity independently varied over time. These light patterns pass through a condenser lens and onto the sample stage to create what we term a ‘light-based augmented reality layer’ ^19^. This can then be used to influence the behaviour of light-responsive microagents that are present. The sample is imaged through a column containing a magnifying tube lens, filters for fluorescent imaging (if required), and finally a microscope objective. Images are captured by a Raspberry Pi 4 with camera module, which performs real-time image analysis. This Raspberry Pi 4 is also used to control a white and UV light emitting diode (LED) for bright field and fluorescence imaging, respectively, and is designed to run analysis code for generating control signals (i.e. the light pattern to be projected). Furthermore, this module is configured as a node in a local *ad-hoc* network to which a Raspberry Pi Zero is also connected. The Raspberry Pi Zero acts as a controller for the projector and is constantly fed information from the imaging process running on the Raspberry Pi 4 through this wireless communication channel. This enables closed-loop control as properties of the system being imaged (e.g. agent location or density) can influence the light pattern being projected. The cost of a DOME varies depending on the configuration, but a system capable of both 9X and 90X magnification and bright-field and fluorescence imaging comes to ~£685 (**Supplementary Table 1**).

A detailed characterisation of the DOME was performed to map its capabilities (**Methods**). Calibrated measurements showed that 1 pixel in the camera field of view (FOV) equated in the real-world to 12 μm^2^ at 9X magnification, and 3.75 μm^2^ at 90X magnification. As the total projection area on the sample stage is 14.5 mm × 26 mm, each projected pixel is ~30 μm^2^ in size. However, due to the sensitivity of the camera and diffusion of the projected light through the optics, a minimum measurable separation between projected and non-projected pixels of 90 μm × 90 μm and 60 μm × 60 μm was found for 9X and 90X magnification, respectively (**Supplementary Figure 1**). We also assessed the light spectrum from the digital light projector and found that at full intensity it contained peaks at 460 nm, 510 nm and 640 nm corresponding to the red, green and blue (RGB) LEDs within the projector (**Supplementary Figure 2**). Little overlap was observed between each of the LEDs’ emission spectra (<2%) making them suitable for multiplexed communication to individual agents (i.e. using each colour as a separate channel). For feedback control, the time taken to image the microsystem, calculate a light pattern, project it and sense this change is crucial. To measure the baseline latency of the DOME, the device was run with no image processing (as this would vary depending upon the experiment performed) to assess the inherent time taken for imaging and projection. We found that the camera resolution was the key determinant of latency, which increased with imaging resolution (**Supplementary Figure 2**). A latency of 0.25 s was found for a typical resolution of 1920 × 1088 pixels.

To show how the DOME can be used to engineer microswarms, we implemented three essential building blocks for collective behaviours using the algae *Volvox* as our microagent. *Volvox* was chosen due to its innate capability to move and sense light ^20^, and because the spherical shape and size of its colonies (350–500 μm in diameter) are easily visualised under a microscope (**Figure 1e**).

First, we focused on the ability to control signalling/communication between agents and shape the spread of information through a population. Signals were encoded as projected light halos around each *Volvox* with a tuneable range and colour (**Figure 2a**). A light signal was transmitted to a nearby *Volvox* if they fell within the halo’s extent. This ‘augmented’ light-based signalling mechanism allowed for a few ‘seed’ *Volvox* colonies that start with the signal active, to propagate the signal throughout the population as they move and interact (**Figure 2b**; **Supplementary Movie 1**) ^21^. The efficiency of signal propagation is governed by the communication range, with larger ranges leading to more rapid spread. In addition, we showed that multiple signals can propagate in parallel (**Figure 2c,d**; **Supplementary Movie 2**) by using different colours for each signal. While the *Volvox* themselves are unaware of the mechanism by which these light signals are transmitted, their motion and interactions play a direct role in the spread of signals and could offer an essential building block in collective behaviours such as consensus formation, quorum sensing, information or disease propagation, or the modelling of extracellular signalling.

**Figure 2:**
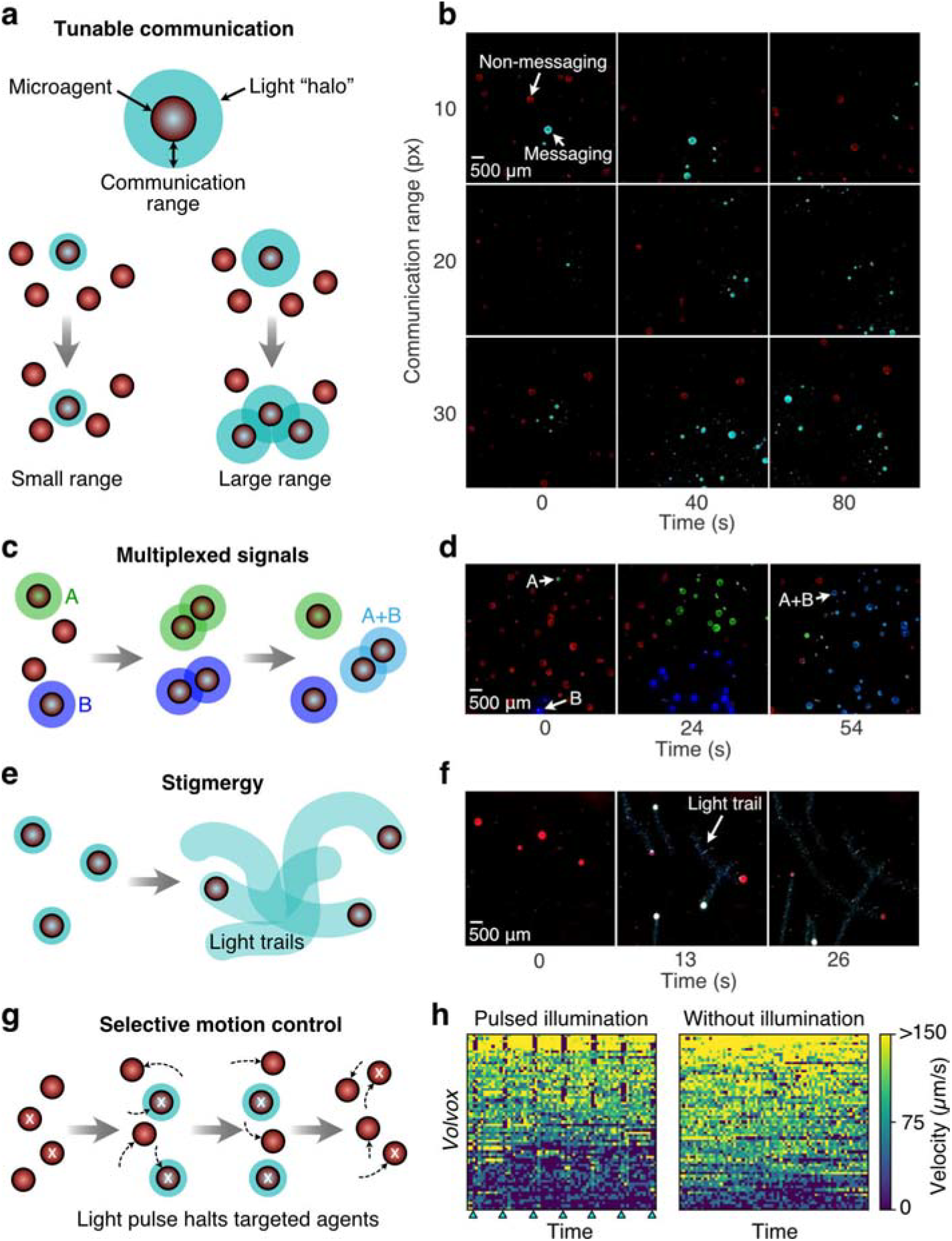
Using the DOME to implement basic building blocks of collective behaviours. (**a**) Schematic of light-based communication between microagents with a tuneable communication range. A larger range increases the probability of a propagation event. (**b**) Image time series of light-based communication between motile *Volvox* colonies with varying communication ranges. Non-messaging *Volvox* appear in red due to illumination by a uniform red background light, while messaging *Volvox* are illuminated in cyan (**Supplementary Movie 1**). (**c**) Schematic of multiplexed communication in which two signals (A: green, and B: blue) propagate between microagents when they are physically close to each other. When the signals interact, they create a third mixed state (A+B: cyan). (**d**) Image time series of multiplexed communication between *Volvox* colonies. Blue and green ‘seed’ colonies are initialised and the two signals propagate through the population until a majority of *Volvox* are in a mixed state (**Supplementary Movie 2**). (**e**) Schematic of light-based stigmergy in which microagents deposit light trails as they move. (**f**) Image time series of light-based stigmergy with *Volvox* colonies depositing cyan light trails (**Supplementary Movie 3**). (**g**) Schematic of selective motion control using the DOME where targeted agents (marked with white crosses) are pulsed with cyan light causing their motion to temporarily halt (**Supplementary Movie 4**). Dotted arrows show the movement of the agents. (**h**) Heat map of experimentally measured *Volvox* colonies over a 17.5 second time course split by (left) colonies exposed to 0.5 second pulses of cyan light (start of time points denoted by triangles), and (right) those without illumination.

Collective behaviours often rely on the ability for agents to modify their local environment through a process called stigmergy, creating a spatially distributed memory that the agents use to coordinate their actions. As a second building block, we implemented light-based stigmergy using the DOME. This was achieved by tracking each *Volvox* colony and projecting light trails over the previous paths they had taken (**Figure 2e**). This resulted in a patchwork of light-trails emerging over time (**Figure 2f**; **Supplementary Movie 3**), reminiscent to the way ants and robot swarms lay trails to optimise foraging or area coverage ^22^.

Serendipitously, while analysing these experiments we observed that many of the moving Volvox slowed down when entering an illuminated path for a short period of time, before resuming their motion (**Supplementary Movie 3**). It is known that *Volvox* are sensitive to light and that they reorient themselves in relation to a light signal due to improve photosynthesis^20^. Therefore, as a final demonstration of a building block of collective behaviours, we attempted to use this innate response as a means to selectively inhibit the movement of a subset of the population. We randomly selected half of the population to be targeted and then periodically after 10 frames (~2.5 s) illuminated these specific colonies for 2 frames (~0.5 s) using cyan light that *Volvox* are known to be responsive to (**Figure 2g**). The velocities for each colony were tracked throughout the experiment. Notably, analysis of *Volvox* motion over time showed that only targeted cells saw clear changes in velocity (slowing down) when illuminated in a punctuated fashion (**Figure 2h**, **Supplementary Movie 4**). However, wide variability in the effect of the light pulses was also observed, with more motile *Volvox* tending to be more strongly inhibited by the light. This heterogeneity is unavoidable in biological systems, further strengthening the need for tailored control of individual agents made possible using the DOME.

In summary, the DOME offers a versatile and low-cost platform for the engineering of microscopic collectives using light. The basic building blocks of local communication, stigmergy, and controllable motion demonstrated could be used as the basis for more complex collective behaviours. Entirely new swarm behaviours could even be engineered by combining the closed loop nature of the DOME with automatic discovery processes based on machine learning algorithms ^23–25^. The DOME’s open-source modular design makes it easy to adapt for new needs, for example, changing the light source of the projector to enable different forms of fluorescent imaging, different magnification, or adding temperature/gas control to maintain the viability of different types of cell (e.g. mammalian cells), and future extensions could even introduce magnetic, sound, or chemical inputs as additional control modalities. Beyond microswarm engineering, the DOME also offers a means to both understand and influence the collective behaviour of natural cellular populations, opening up new avenues for the study of complex systems spanning cancer to the microbiome.

## Methods

### Fabrication of the DOME

The DOME is an assembly of parts 3D printed in PLA plastic. The original device was printed using an Ultimaker 2+ printer, and has been replicated using an Anycubic i3 Mega, a lower specification machine. Parts were designed using Autodesk Inventor and Fusion 360 and can be printed without requiring dissolvable supports. *Z*-plane focusing is achieved using a linear rail set, where manually rotating the lead screw raises or lowers the sample stage with respect to the imaging lens and camera. An *x*-*y* translational stage is attached to the sample stage to allow easy sample adjustment in this plane. This affects only the positioning of the sample and has no bearing on the relative positions of the optical components. The imaging column faces upwards towards the sample stage from the base of the DOME and contains an optical filter holder on a pivot hinge and printed internal threads to attach magnification lenses. The imaging column has two different attachments that can be used, one for a lower magnification option in which only a tube lens is required and an extended version for higher magnification, where a microscope objective can be attached.

### Optical set up

The digital light projector (DLP) module is fixed on the sample stage, and thus is unaffected by any adjustment in *z*-plane focus. It is instead focused independently using a screw lever attached to the projector. Light from the projector is focused by a condenser lens (50mm diameter PCX condenser lens, Edmund Optics), resulting in a total projection size of 14.5 mm × 26 mm. A white LED (RS Components) can also be attached behind the condenser lens to act as a brightfield light for standard microscopy. To provide an illumination across the sample, a ground glass diffuser (Thor Labs) is placed between the LED and condenser lens. Note that although the DOME is capable of bright field illumination, this feature was not used in the experiment presented here. A camera (Camera Module V2, Raspberry Pi) sits on the base of the imaging column pointed upwards at the sample stage. Optics such as wavelength or neutral density filters can be added into the optics holder within the imaging column on an application specific basis. For the lower magnification configuration, the imaging column ends with a 9X tube lens (Eyepiece Cell Assembly, Edmund Optics) screwed into a threaded cylindrical casing. For higher magnification applications the cylindrical section is extended, ending in an RMS thread to fit a standard microscope objective (10X Semi-Plan Standard Objective, Edmund Optics). While the length of this lens piece is specific to the lenses used here, due to the modular nature of the DOME it would be trivial to adjust this dimension to suit alternative optics. Positioning both the camera and projector perpendicular to the sample stage results in significant lens flare through which imaging is difficult. To circumnavigate this the projector is angled at 10°, positioning the bright spot created by the light source of the projector out of the camera field of view (FOV).

### Characterisation of imaging and projection modules

The modular design of the DOME allows for interchangeable levels of magnification using a tube lens and an RMS threaded tube to mount different microscope objectives. A low magnification of 9X is suitable for larger microsystems of the order of hundreds of microns in size such as multi-cellular algae, while a higher magnification of 90X is appropriate for smaller agents such as mammalian cells or bacteria.

To assess the imaging and projection capabilities, we began by comparing the 9X and 90X magnification settings and used *Volvox* as an example subject. *Volvox* are an algae 350–500 μm in diameter where a single spherical colony houses up to 50,000 cells. At 9X magnification, many colonies can be seen in low detail whereas at 90X magnification only a few at are visible but smaller features such as daughter colonies within the body of each *Volvox* are clearly seen (**Figure 1e**). A scale for both magnifications was calculated by imaging a measuring ruler.

Next, we considered light projection. When light from the projector is focused through the condenser lens, the total projection area on the sample stage is 14.5 mm × 26 mm, making each projected pixel theoretically 30 μm × 30 μm in size. Due to differing FOVs for each magnification, this fixed projection area leads to a trade-off with the number of projected pixels that are visible to the camera (300 × 300 pixels for 9X, and 88 × 66 pixels for 90X magnification).

To test the precision of projected light patterns, a series of line triplets of differing size and spacing were projected onto a neutral density filter (**Supplementary Figure 1a,b**). The resulting camera images were then analysed by averaging the intensity for each pixel row (**Supplementary Figure 1c,d**). High precision projected patterns would result in clear differences in light intensity for even closely spaced lines. For 9X magnification, lines of 1-pixel width (30 μm) were difficult to distinguish, with improvements at separations of 2 pixels and clear differences at separations of more than 3 pixels (90 μm). For the 90X magnification, distinct peaks are seen for lines separated by just 2 pixels (60 μm). Measuring across the peaks for each line set in pixel distance and multiplying by the scale factor allows direct measurement of the observed projector pixel size. At both magnifications this agreed with the theoretical projector pixel size of 30 μm × 30 μm.

### Light spectra measurements

Another key feature of the DOME is the ability for each projected pixel to have a different colour. This offers the means to provide multiple light-signals to different agents and supports multiplexed communications for more complex behavioural control. This capability is possible due to the projector containing three separate LEDs for red, green and blue light. As living microagents are often sensitive to limited wavelengths of light, we characterised the light spectra of each LED separately. The light spectra produced by the projection module was performed using a calibrated spectrophotometer (Ocean Optics). To collect the readings the optical fibre used for measurement was attached to the DOME at the sample plane facing upwards. The projector was set to a full screen display where all pixels had value (0, 0, 255), (0, 255, 0) or (255, 0, 0) respectively for red, blue and green.

### Latency of closed-loop control schemes

For effective closed-loop control, the latency between the imaging of the sample and resulting light projection should be minimised. For the DOME, latency can be described as the time period between subsequent camera frames being obtained, since a new frame is captured only after the imaging module has sent data to the projection module and received confirmation of its receipt. The system latency will vary between applications and depends on camera setting, and the extent of image processing required per frame. To characterise the baseline latency of the DOME, an experiment was run with no image processing where the projector was switched between an on (white) and off (black) state after every camera frame received. We found that the primary source of latency was the time needed by the camera to capture an image, which is dependent on the resolution used. The algorithm was therefore run over a range of resolution settings. Each time measurement was taken as a mean average over 100 frames, running at a shutter speed of 100 milliseconds. Measurements were taken at 63 different resolution settings, starting at 640 × 480 pixels and increasing in increments of 32 × 32 pixels until maximum camera resolution was reached at 2646 × 2464 pixels. The results showed that even for the highest resolutions, we are able to close the loop in under one second (**Supplementary Figure 3**). This means that, assuming a single-agent projection area of 2 × 2 pixels (60 μm × 60 μm), any agent moving slower than 83 μm/s could be imaged at the highest possible resolution (latency of 0.725 s) whilst maintaining accurate light projection in relation to their position. More typically, the DOME is operated at a resolution of 1920 × 1088 pixels for which the latency is 0.25 s. At this resolution, agents moving slower than 240 μm/s can be accurately tracked by projected light. For faster moving agents, lower projector resolutions can be employed, larger light projection areas used, or both. More advanced algorithms can also be used to predict the location of an agent in a subsequent frame based on its current trajectory, however, this has not been necessary for the applications we have explored.

### Closed-loop computational set up

The projection module is comprised of the DLP LightCrafter Display 2000 Evaluation Module (Texas Instruments) interfaced through a custom PCB (Pi Zero W adapter board, Tindie) with a Raspberry Pi Zero W. The imaging module comprises a Raspberry Pi 4, Raspberry Pi camera and illumination LEDs. This computer acts as the primary computing module and user interface, and can be connected to a monitor, mouse and keyboard, or accessed remotely. Crucial to the closed-loop control scheme of the DOME is two-way communication between the imaging and projection module. Due to the interface between the Raspberry Pi Zero and DLP unit there are no ports available to facilitate a physical connection. As an alternative, the two Raspberry Pi modules are configured as nodes in an *ad-hoc* wireless network. The network was established by editing the network interface files on both Raspberry Pi computers to include details of the required *ad-hoc* connection and IP addresses for both nodes. This ad hoc configuration allows the two-way transfer of information for closed-loop control, with the imaging module operated as a server socket, and the projection module connecting as a client. The connection also enables the user to control the projection module from the imaging module via a VNC connection. With the projection module Pi running VNC server and the imaging module Pi running VNC Viewer, both desktops can be accessed and controlled using a single desktop, mouse and keyboard set up if needed.

### Calibration algorithm for the camera and projector

Due to the nature of imaging through a circular imaging column and lenses, raw camera images contain sizable “dead space” (area that is imaged but for which the projectors light pattern does not extend too). A typical raw camera frame will appear as a black rectangle with a circular area in the centre in which the sample is visible. To increase image processing efficiency and reduce file sizes, the first step in the calibration process is to crop the total FOV down to a rectangular area that fits inside the circular area of visibility. For this, contour detection is run to find the illuminated area. From these coordinates, the largest square is found by contour detection and this information is written to a file in the format (centre-x, centre-y, width, height). This file can then be imported by all other programs to maintain consistency. Critical to the operation of the DOME is the ability to translate coordinates within the camera frame of reference into the corresponding projector coordinates. For this, the camera space is mapped to the projector space through a calibration process. The first step is to locate approximately where in the projector space the camera is focused using an iterative quadrant search. Once the appropriate sub-space has been found, a 4-point square is projected into this area and located in the camera frame using contour detection. With these sets of coordinates for the projector and camera spaces, the parameters for a matrix transformation operation can be extracted. The baseline code for calibration is provided as part of the open-source DOME software (see Data Availability section).

### Agent imaging and tracking

The Raspberry Pi Camera was used for all imaging, operating at a resolution of 1920 × 1088 pixels. In all cases, exposure mode was set to ‘spotlight’ and camera ISO was set to 100, with a shutter speed of 200 ms. The camera was operated using the capture continuous method in which images are captured in an infinite loop, iterating over frames. A neutral density filter (NE510B-A, Thor Labs) was placed into this holder to minimise optical interference artifacts. All image processing and projection algorithms were run on Python through the Raspberry Pi OS. *Volvox* agents were detected in the camera FOV by finding image contours using OpenCV and filtering for size and compactness. ID based tracking was implemented by matching the locations of contours in a given frame to those in the previous frame. Agents were matched to their most likely ID by checking their current location against agents in the previous frame. The closest match was assumed to be the same agent, provided that the distance between the two locations was smaller than 35 pixels (420 μm). This method was largely effective in locating and matching agents. However, we found it did not always reliably distinguish agents where two or more collided. This issue was, for the most part, minimised by the relatively low density of *Volvox* in the FOV at any one time. Where this issue did occur, the tracking system is designed to assign new IDs to the agents once separated to avoid data becoming biased by these events.

### Agent illumination

*Volvox* samples (Blades Biological, UK) were maintained at room temperature. For imaging, 75 μL of the *Volvox* suspension was added to the sample arena, comprised of 3D printed walls attached to a microscope slide (**Supplementary Note 1**). As the *Volvox* are placed into a confined arena with depth multiple times larger than their diameter they are able to move freely. During experiments with *Volvox*, samples were uniformly illuminated with a low-level red light. Due to the off-axis projection, this produced images of *Volvox* agents that appear bright red against a dark background (**Figure 2**), as opposed to bright-field imaging in which agents are dark against a brightly lit background (**Figure 1e**). Keeping background light levels low in this way, and using red light minimises the effect of background light on *Volvox* agents, which respond much more strongly around the 500 nm wavelength range. A typical projection image was a uniformly dark red background, RGB pixel values (50, 0, 0), with coloured patterns at a brighter intensity. Light patterns were generated based on agent location coordinates, sent as a JSON formatted data to the projection module after translated from camera to projector space.

## Supporting information

Supplementary Movie 1 - Communication Volvox

Supplementary Movie 2 - Dual Communication Volvox

Supplementary Movie 3 - Stigmergy Volvox

Supplementary Movie 4 - Motion Control Volvox

## Data availability

All part designs and supporting software required to create a working DOME are available through a public Bitbucket repository: http://bitbucket.org/hauertlab/dome

## Acknowledgements

This work was supported by an EPSRC DTP scholarship (A.R.D), the European Union’s Horizon 2020 FET Open programme under grant agreement. No. 800983 (S.H), BrisSynBio, a BBSRC/EPSRC Synthetic Biology Research Centre grant BB/L01386X/1 (S.H., T.E.G.), and a Royal Society University Research Fellowship UF160357 (T.E.G.)

