## Supplementary Information for "An open platform for high-resolution light-based control of microscopic collectives"

| **Supplementary Notes** | **Page** |
|---|---|
| Supplementary Note 1: Arena for *Volvox* experiments | 2 |
| **Supplementary Figures** | |
| Supplementary Figure 1: Characterization of projection and imaging modules | 3 |
| Supplementary Figure 2: Light spectra of projection module | 4 |
| Supplementary Figure 3: Latency of closed-loop control applications | 5 |
| **Supplementary Tables** | |
| Supplementary Table 1: Breakdown of DOME component costs | 6 |
| **Supplementary Movie Captions** | |
| Supplementary Movie 1: Communication | 7 |
| Supplementary Movie 2: Programmable messaging | 7 |
| Supplementary Movie 3: Stigmergy | 7 |
| Supplementary Movie 4: Motion control | 7 |

### Supplementary Note 1: Arena for *Volvox* experiments

To allow the free movement of *Volvox* colonies, a sample area (see image below) was constructed. A square chip of outer dimensions 25 mm x 25 mm with a 7.75 mm x 7.75 mm square cut from the middle was 3D printed in PLA and attached to a standard glass microscope slide using superglue adhesive. This square well has depth of 1.5 mm, allowing *Volvox* colonies of 350–500 μm in diameter to move freely in the *x*-*y* plane, with some limited movement in the *z* plane.

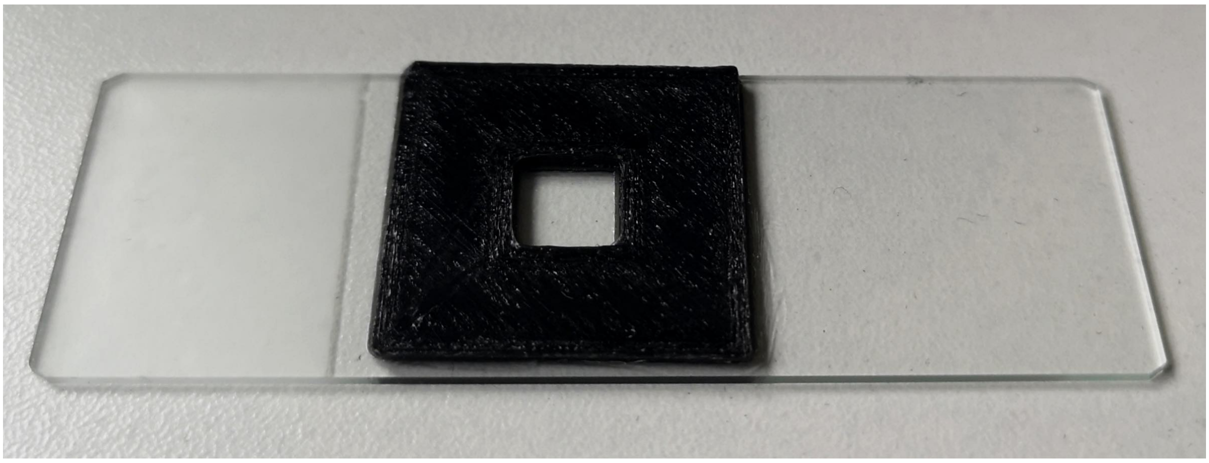

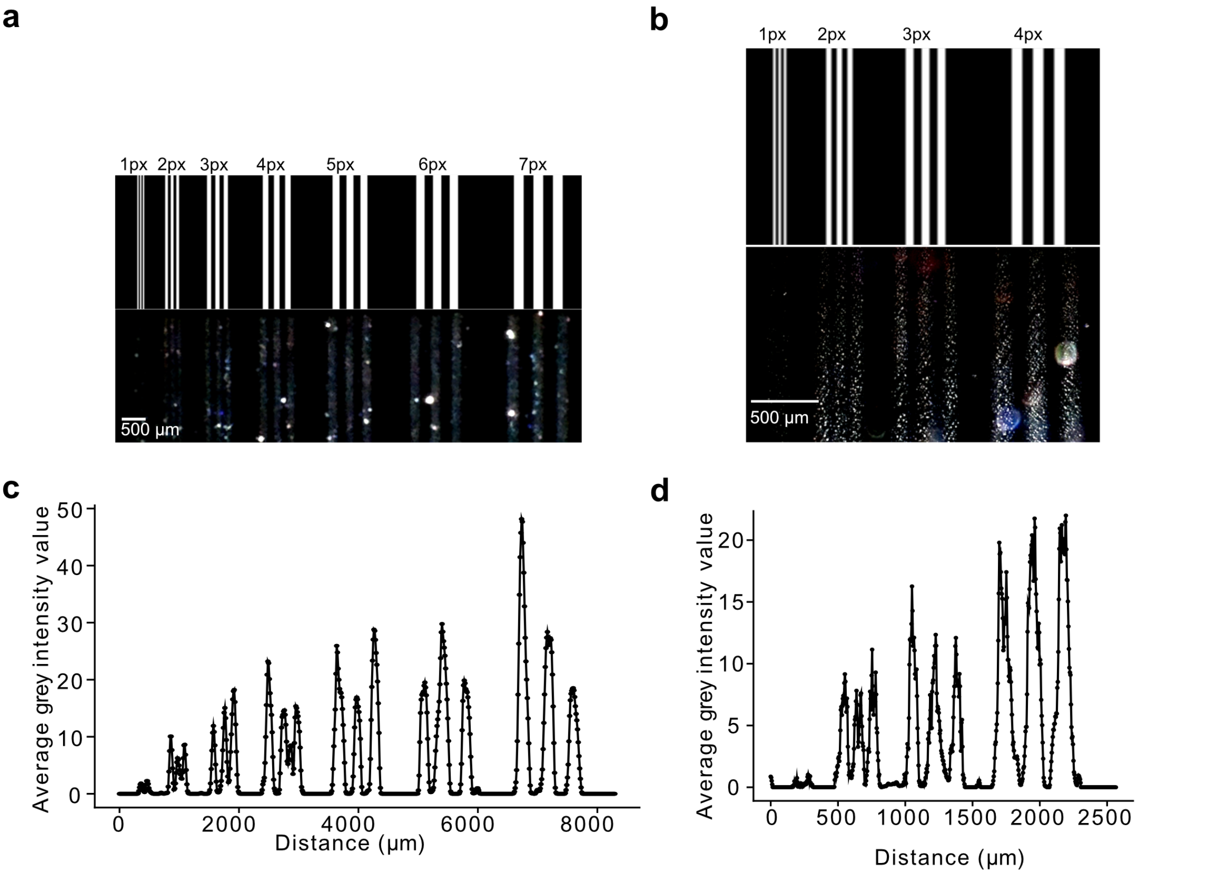

**Supplementary Figure 1: Characterisation of projection and imaging modules.** (**a**) Projection image of line triplets of increasing width up to 7 pixels for 9X magnification (upper) and corresponding camera image (lower). (**b**) Projection image of line triplets of increasing width up to 4 pixels for 90X magnification (upper) and corresponding camera image (lower). (**c**) Intensity plot across test image for 9X magnification, measured as the average grey-scale value for each pixel column in the image. (**d**) Intensity plot across test image for 90X magnification, measured as the average grey-scale value for each pixel column in the image.

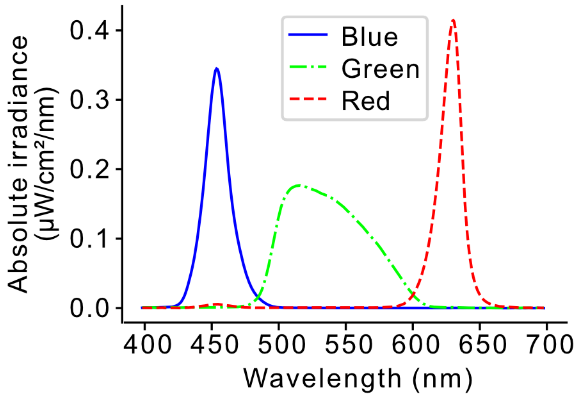

**Supplementary Figure 2: Light spectra of projection module.** Spectra of light emission from the projection module showing clear separation of the red, green and blue LEDs.

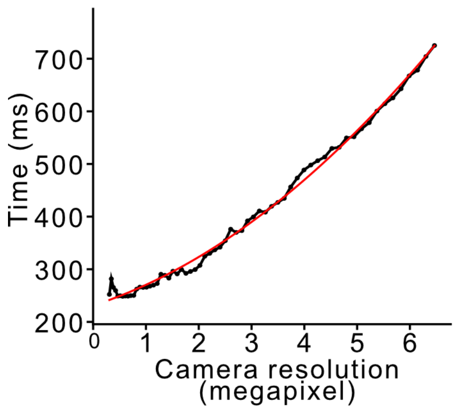

**Supplementary Figure 3: Latency of closed-loop control applications.** Latency of closed-loop control as a function of camera resolution. Red line shows a second order polynomial fit of $y = 6.8x^2 + 32.6x + 231$.

**Supplementary Table 1: Breakdown of DOME component costs.**

| *Optical* | **Cost (£)**[a] |
|---|---|
| Projector (DLP Lightcrafter Display 2000 EV, Texas Instruments) | 109 |
| Condenser lens (50mm Diameter PCX , Edmund Optics) | 37 |
| Tube lens (9X Eyepiece Cell Assembly, Edmund Optics) | 61 |
| 10X objective (Semi-Plan Standard Objective, Edmund Optics) | 122 |
| Glass diffuser (DG10-1500, Thor Labs) | 15 |
| Neutral density filter (NE10B-A, Thor Labs) | 47 |
| Longpass filter (FEL0500, Thor Labs) | 60 |
| *Electrical* | |
| Raspberry Pi (Raspberry Pi 4 Model 4GB, The Pi Hut) | 54 |
| Raspberry Pi (Raspberry Pi Zero W, The Pi Hut) | 9 |
| Camera (Raspberry Pi Camera V2, The Pi Hut) | 24 |
| 2 × SD card (SanDisk Ultra 16GB microSDHC, Amazon) | 14 |
| Interface PCB (Pi Zero W DLP2000EVM adaptor board, Tindie) | 3 |
| Power supply (UK Raspberry Pi 4 Power Supply The Pi Hut) | 8 |
| Power supply (Raspberry Pi 3 Universal Power Supply) | 8 |
| *Mechanical* | |
| PLA filament (Black Premium PLA 1.75mm, FilaPrint) | 28 |
| Linear rail set (Glvanc 3D Printer Guide Rail Sets, Amazon) | 21 |
| *x-y* stage (Zetiling Microscope Moveable Stage, Amazon) | 15 |
| Linear Motion Ball Bearing (LM8LUU, Amazon) | 7 |
| Lighting and fastening sundries | 41 |
| **Total cost** | **685** |

a. All prices given to the nearest pound (£) and inclusive of 20% VAT.

### Supplementary Movie Captions

**Supplementary Movie 1: Communication.** Light-based communication between motile *Volvox* colonies with a 20 px communication range. Non-messaging *Volvox* appear in red due to illumination by a uniform red background light, while messaging *Volvox* are illuminated in cyan. Propagation of the message can be seen throughout the population over time.

**Supplementary Movie 2: Programmable messaging.** Programmable messaging between *Volvox* colonies. A blue and green seed are initialized and propagate through the population until a majority in the mixed state colored in cyan is reached.

**Supplementary Movie 3: Stigmergy.** Light-based stigmergy with *Volvox* colonies depositing light trails in cyan. Agents can be seen to change velocity upon encountering a light trail.

**Supplementary Movie 4: Motion control.** Selective light-based control of *Volvox* movement. Half the *Volvox* population (randomly selected) are illuminated with blue light for 2 frames every 10 frames, causing a slowing of their movement.
